# Survival and detection of bivalve transmissible neoplasia from the soft-shell clam *Mya arenaria* (MarBTN) in seawater

**DOI:** 10.1101/2021.12.25.473725

**Authors:** Rachael M. Giersch, Samuel F.M. Hart, Satyatejas G. Reddy, Marisa A. Yonemitsu, María J. Orellana Rosales, Madelyn Korn, Brook M. Geleta, Peter D. Countway, José A. Fernández Robledo, Michael J. Metzger

## Abstract

Many pathogens can cause cancer, but cancer itself does not normally act as an infectious agent. However, transmissible cancers have been found in a few cases in nature: in Tasmanian devils, dogs, and several bivalve species. The transmissible cancers in dogs and devils are known to spread through direct physical contact, but the exact route of transmission of bivalve transmissible neoplasia (BTN) has not yet been confirmed. It has been hypothesized that cancer cells could be released by diseased animals and spread through the water column to infect/engraft into other animals. To test the feasibility of this proposed mechanism of transmission, we tested the ability of BTN cells from the soft-shell clam (*Mya arenaria* BTN, or MarBTN) to survive in artificial seawater. We found that BTN cells are highly sensitive to salinity, with acute toxicity at salinity levels lower than those found in their environment. BTN cells also survive longer at lower temperatures, with >48% of cells surviving a week in seawater at temperatures from 4°C to 16°C, and 49% surviving for more than two weeks at 4°C. With one clam donor, living cells were observed for more than eight weeks at 4°C. We also used qPCR of environmental DNA (eDNA) to detect the presence of BTN-specific DNA in the environment. We observed release of BTN-specific DNA into the water of aquaria from tanks with highly BTN-positive clams, and we detected BTN-specific DNA in seawater samples collected from BTN-endemic areas, although the level detected was much lower. Overall, these data show that BTN cells can survive well in seawater, and they are released into the water by diseased animals, supporting the hypothesis that BTN is spread from animal-to-animal by cells through seawater.

## INTRODUCTION

Most cancer stays with the organism from which it came, arising and dying within a single host, but in a few cases, cancer has evolved to transmit from one animal to the next, acting as a pathogen as well as a cancer. The first naturally transmissible cancers to be found were the canine transmissible venereal tumor (CTVT) (1, 2) and the Tasmanian devil facial tumor disease (DFTD) (3, 4). More recently, a leukemia-like disease in multiple bivalve species, called disseminated neoplasia (DN) or hemic neoplasia, was shown to be a transmissible cancer (5). DN is characterized by proliferation of non-adherent, rounded, polyploid neoplastic cells primarily found in the hemolymph of infected bivalves, which disseminate into tissues during later stages of this typically fatal disease (6, 7). The disease had been reported for many decades, although the etiology was unknown. Retroviruses or pollution had been thought to be the likely causes (8, 9), although two early reports had suggested that it could be due to infectious spread of cancer cells (Sunila et al. 1998 and James Moore’s 1993 Ph.D. Dissertation (10, 11)). DN was first confirmed to be a transmissible cancer in soft-shell clams (*Mya arenaria*) (5), and later, multiple independent lineages of transmissible cancer were identified in multiple species worldwide (12). To date, seven lineages of the bivalve transmissible neoplasia (BTN) in eight bivalve species have been reported (5, 12–17). In many cases, the BTN that circulates in each species has arisen from a member of that same species. However, increasing cases of cross-species transmission have been reported (a lineage from *Venerupis corrugata* to *Polititapes aureus* and a single lineage of *Mytilus* BTN now found in four *Mytilus* species). Furthermore, multiple independent lineages of BTN have been found to circulate within the same species (as in *Cerastoderma edule* and *Mytilus trossulus*). In soft-shell clams, all analyzed samples of DN since the confirmation of cancer transmission so far have proven to be a part of one lineage of BTN that arose from a single founder soft-shell clam and has since spread throughout populations along the North American East Coast between Prince Edward Island (Canada) and New York (USA). DN has been reported in *M. arenaria* as early as 1978 (18, 19) and it has been observed as far south as Chesapeake Bay, Maryland, USA (9, 20, 21). While it is likely that these earlier reports of DN in soft-shell clams represent the same BTN lineage observed today, this cannot yet be confirmed.

CTVT and DFTD are well known to be transmitted by close physical contact (during sex and biting, respectively), but most adult bivalves are sessile or with limited mobility, without direct contact of soft body tissues, making widespread BTN transmission through direct contact unlikely. The genomic evidence clearly shows that cancers in different individual bivalves do not match the genotypes of their hosts and that cancers in different individuals, separated by large distances (and in some cases in different species and different oceans), are nearly identical, so the cancer lineages must transmit through some alternate mechanism. The most likely route of transmission is release of the BTN cells into the seawater and uptake of the cells by naïve animals through filter feeding. For this to occur, BTN cells need to be released from animals and survive in seawater long enough to engraft into a naïve individual.

A previous report (the first known to propose a transmissible cancer hypothesis for DN in bivalves) characterized the survival of cancer cells from soft-shell clams collected from Chesapeake Bay in 1989 and showed that they survive well for 6 hours, and that survival can be affected by salinity and temperature (10). As in Sunila et al., we aim to analyze the ability of BTN cells from soft-shell clams (*Mya arenaria* BTN, or MarBTN) to survive in seawater. We determined the effect of salinity, pH, and temperature on their short-term survival in artificial seawater (ASW), and we determined how long the cells can survive at varying temperatures. We additionally tested whether BTN cells can be detected in seawater through the analysis of environmental DNA (eDNA) collected from both laboratory and field settings. Our findings provide further validation supporting seawater-based transmission of BTN in the wild.

## MATERIALS AND METHODS

### Collection of clams and MarBTN cells

Soft-shell clams (*M. arenaria*) were collected by commercial sources from multiple locations in Maine, USA (Table S1), and animals were screened for high levels of DN as before (5). Animals were housed in 1× ASW (36 g / L Instant Ocean, Blacksburg, VA, USA), in aerated aquaria, supplemented 2-3 times weekly with PhytoFeast or LPB Frozen Shellfish Diet (Reed Mariculture, Campbell, CA, USA). Approximately 0.5-1 mL of hemolymph was collected from the pericardial sinus of each animal using a 0.5 in 26-gauge needle on a 3 mL syringe. Several drops were placed in a well of a 96-well plate and incubated at 4-10°C for 1 hour to allow the cells to settle. Wells were screened for clams with high levels of BTN based on morphological differences between healthy hemocytes and BTN cells on an inverted phase-contrast microscope. Only animals with ≥75% of cancer cells in their hemolymph were used in survival experiments.

### Counting live cells in artificial seawater

We used the alternate vital dye, erythrosine B (MilliporeSigma, Burlington, MA, USA) to stain samples to discriminate live and dead cells during counting. This dye is soluble throughout all salinities used in this study. To count live cells, 10 μL of ASW containing cells were mixed 1:1 with 2× erythrosine B solution (10 μg/mL, dissolved in PBS4, which is PBS plus 400 mM NaCl to approximate marine salinity). After 10 min at room temperature, live cells were counted manually on a hemocytometer, counting only rounded, refractive cells that exclude dye.

### Short-term cell survival assays

For short-term MarBTN cell survival assays, hemolymph was collected from heavily neoplastic animals, and allowed to sit on a 6 cm tissue culture dish or 24-well plate at 4 °C for one hour to allow any healthy hemocytes to adhere to the dish so that they could be removed. Non-adherent cells were then removed and spun down at 500 × g for 10 min at 4 °C. Hemolymph was removed, and cells were resuspended in 1× ASW: filter-sterilized Instant Ocean with no additives, 36 g/L, specific gravity (sg) 1.023, and pH 7.93. For salinity, cells were diluted to an approximate concentration of 1 × 10^6^ cells/mL, and 20 μL of cells were added to 180 μL of ASW with varying concentrations of Instant Ocean, from 0 to 2× the normal salinity level (with sg measured by a refractometer) and placed in wells of a 96-well plate at 16°C. Cell counts at 4 hours were normalized to expected concentration calculated pre-dilution, as cell death in low salt was too rapid to allow for counting after dilution. For pH, cells were aliquoted into one tube for each condition, spun a second time, and resuspended in 200 μL of ASW with different pH (3.8-9.3, modified with 2M NaOH or 3N HCl), with a target of 1 × 10^6^ cell/mL, and put into wells of a 96-well plate. For temperature, the cells were resuspended in 400 μL of 1× ASW (sg 1.023, pH 7.93) and placed in wells of a 24-well plate at the indicated temperatures (4-37°C). For pH and temperature, cell counts at time zero were used to normalize cell survival.

### Long-term cell survival assays

For long-term cell survival, penicillin/streptomycin (1×, GenClone, Genesee Scientific, El Cahon, CA, USA) and voriconazole (1 mM final concentration, Acros Organics, Thermo Fisher Scientific, Waltham, MA, USA) was added to ASW. Other antimicrobial drugs were tested (e.g., moxifloxacin, doxycycline, metronidazole, and triclosan) but were not found to reduce contaminants at a concentration that was non-toxic to BTN cells. Cells were collected as above and resuspended in 400 μL 1× ASW with penicillin/streptomycin/voriconazole, in wells of a 48-well plate, at 2 × 10^5^ – 2 × 10^6^ cells/mL. After each timepoint, the volume was measured by pipette, and ASW with antimicrobial drugs was added to replace the media removed for cell counting and lost due to evaporation. Live cell counts at each timepoint were normalized by live cell counts at time zero to calculate survival, and the normalization value was multiplied by 0.975 after each additional timepoint to reflect the removal of 10 μL of cells from the original 400 μL sample.

### eDNA extraction from aquaria

Animals were maintained at approximately 10°C in individual tanks in 1 L of 1× ASW, with constant aeration. 24 hrs prior to water collection, the entire volume of the tank water was replaced. Each day for three days, 250 mL of each water sample was collected, and the entire tank water was replaced, so that each sampling is from water with 24 hrs of exposure to a single clam. Water samples were vacuum filtered through a 47 mm 0.45 μm cellulose nitrate filter. Using forceps, the filtered sample/paper was folded small enough to fit into a 2 mL tube and frozen at −80°C until extraction was performed.

Extraction protocol was modified from Renshaw et al. (22). Briefly, 900 μL CTAB buffer (2% CTAB w/v, 20 mM EDTA, 100 mM Tris-HCl, and 1.4 M NaCl, in water) was added to the filter, and the tubes were incubated at 65°C for 30 min. Tubes were spun to collect the sample in the bottom of the tube and 900 μL chloroform:isoamyl alcohol (24:1) was added, followed by shaking or vortexing. Tubes were spun for 5 min at 15,000 × g, and the 700-850 μL aqueous layer was transferred to a new tube with 700 μL chloroform. This was shaken and spun as before and the ~700 μL aqueous layer was transferred to a new tube with 700 μL cold isopropanol and 24 μL 5M NaCl. 4.67 μL glycogen blue was added to ensure visibility of the pellet, and samples were allowed to precipitate overnight at −20 to −30°C. DNA was spun for 10 min at 15,000 × g and the liquid removed by pipette. 500 μL of 75% ethanol was slowly added and poured off. DNA pellets were air dried and resuspended in 100 μL Buffer EB (Qiagen, Hilden, Germany).

### Seawater collection and extraction of MarBTN eDNA

Seawater samples were collected from surface water overlying clam-flats in Maine by filling a single 4-liter acid-washed (5% HCl) HDPE bottle from each location (Quahog Bay Dam, June 6, 2021, 43.812541, −69.896802; Gurnet Landing, June 6, 2021, 43.853734, −69.898677; and Long Cove, June 13, 2021, 43.777156, −69.958582). Sample bottles were stored in a cooler with ice packs until delivery to Bigelow Laboratory within 24 hours of collection. Triplicate sub-samples of 500 mL seawater were filtered from each bottle onto 47 mm diameter, 0.2 μm Supor filters (Pall Corp., Ann Arbor, MI, USA) to collect environmental DNA. Filters were rolled and placed in 4.5 mL cryovials (USA Scientific, Ocala, FL, USA) for storage at −80°C until DNA extraction. Filters were rolled to ensure that the particle-bearing filter surface faced inward and that the filter would unfurl when it was transferred to a DNA extraction tube.

Environmental DNA was extracted from the Supor filters using the DNeasy PowerWater kit (Qiagen). Frozen filters were transferred from the cryovials to the 5 mL PowerWater bead tubes and 1 mL of warmed (55°C) PW1 solution was added. Bead tubes containing a filter, PW1 solution and garnet beads were vortexed for 30 min on a Vortex Genie IIT (Scientific Industries, Bohemia, NY, USA) using a 15 mL tube adapter. After the bead-beating step, the crude cell lysate, extracted Supor filter, and most of the beads were tapped into the barrel of a sterile 10 mL syringe held over a 2 mL Eppendorf DNA LoBind tube to catch sample lysate. The syringe’s plunger was inserted a short way into the syringe barrel before the syringe assembly was flipped upright to purge air. The syringe assembly was inverted over the 2 mL LoBind tube for a second time, and the remaining lysate was pressed out of the bead and filter slurry. The volume of this crude sample lysate was recorded, and the remainder of the DNA extraction procedure followed the kit protocol. Extracted DNA samples were stored in DNA LoBind tubes at −20°C until analysis by qPCR.

### qPCR of hemocyte DNA and eDNA

To quantify the presence of neoplastic DNA in a hemolymph genomic DNA or eDNA sample, allele-specific qPCR was performed using four sets of primers (Table S2). The primary locus was a MarBTN-specific insertion of the LTR-retrotransposon *Steamer* at the *N1N2* gene. A MarBTN-specific primer pair targeting this insertion junction amplifies ½ the total amount of *N1N1* alleles in a cancer cell (as the insertion is in two of four copies of the gene in a tetraploid region) and a primer pair in a conserved region of the *N1N2* ORF nearby quantifies the total copies of the *N1N2* locus present. The ratio of the two can be used to determine the fraction of clam hemolymph made up of MarBTN cells. A single plasmid (pCR-SteamerLTR-N1N2) was used for the standard curve. It was made by cloning the *Steamer*-N1N2 amplicon, amplified from genomic DNA of MarBTN cells (Zero Blunt TOPO PCR cloning kit, Invitrogen, Waltham, MA). The secondary marker was a different MarBTN-specific *Steamer* integration site, termed HL03 (23). A plasmid was cloned which includes both the HL03 locus and a separate conserved region of the *EF1α* gene as a control (pIMHL03c2-EF1α). Primers used for cloning control plasmids are listed in Table S2, and sequences have been submitted to GenBank. The plasmid concentration was measured (Qubit, Thermo Fisher Scientific) and copy number per μL was calculated based on the plasmid sizes. Plasmids were linearized with 0.25 μL of NotI-HF (NEB, Ipswitch, MA, USA) for 30 min at 37°C in a 20 μL reaction at 1 × 10^10^ copies/μL, heat-inactivated 20 min at 65°C, then diluted to 1 × 10^9^ with 180 μL Buffer AE (Qiagen). Standard curves were prepared from 1 × 10^7^ copies/rxn to 1 × 10^1^ copies/rxn. For aquaria samples, 2 μL of extracted eDNA was run in 10 μL reactions on a StepOnePlus real-time PCR cycler (Applied Biosystems, Waltham, MA, USA). For field eDNA samples, 4 μL of eDNA was used in a 20 μL reaction for increased sensitivity. Reactions were run as follows: 95°C for 2 min, 40 cycles of 95°C for 15 s and 60°C for 30 s, followed by a melt curve using 95°C for 15 s, 60°C for 1 min, and ramping 0.3°C from 60°C to 95°C, followed by a 15 s hold at 95°C. All samples were run in triplicate and values presented are an average of triplicates, treating wells with undetectable amplification as zero copies.

## RESULTS

In order to determine the factors that affect survival of *M. arenaria* BTN cells in seawater, we collected MarBTN cells from heavily neoplastic animals and incubated those cells in ASW of varying salinity, pH, and temperature. Identification of dead cells is challenging in marine cells, as trypan blue (a vital exclusion dye often used to identify dead cells in mammalian cell culture) precipitates out of solution when prepared at salinities found in seawater. Therefore, we tested alternate vital dyes, and found that erythrosine B remains in solution and functions well at salinities up at least twice marine salinity (72 g/L Instant Ocean, 1.045 sg).

As found in the previous study of bivalve DN cells, MarBTN cells rapidly die in low salinity water (Figure 1A). In contrast, the majority of cells survive at least 4 hrs in ASW of expected marine salinity in the New England area (1.023 sg). BTN cells also show the greatest survival at expected marine pH, but complete cell death required highly acidic conditions not likely to be relevant to the environment (Figure 1B). Variation of temperature from 4 to 37°C, in contrast, had minimal effect on survival within 4 hours (Figure 1C).

**Figure 1.**
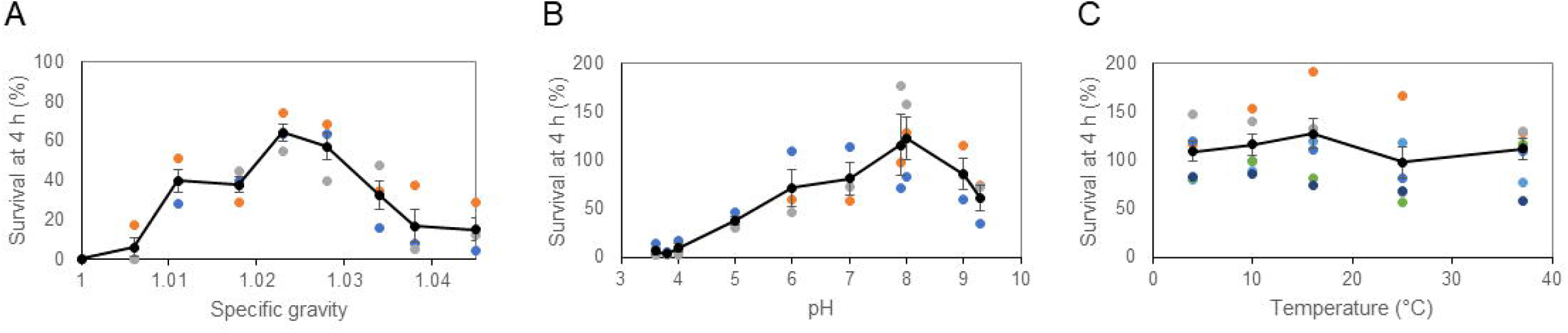
The effect of salinity, pH, and temperature on survival of MarBTN cells in artificial seawater. BTN cells from soft-shell clams were collected and incubated in ASW with varying (***A***) salinity, (***B***) pH, and (***C***) temperature for four hours before survival was measured, using erythrosine B to identify viable cells. Unless it was the variable being tested, ASW was prepared at average marine salinity (1.023 sg), without additional pH modification (pH 7.93), and held at 16°C. For pH and temperature experiments, live cell counts at 4 hrs were normalized by cell counts for each well at initiation of the experiment, but due to acute toxicity of low salinity, this could not be done for part A. These counts were normalized by the expected number of cells based on cell counting of the initial cell suspension, assuming no loss during centrifugation and pipetting. For each experiment, 3-6 independent replicates using BTN cells from separate diseased clam donors were conducted (colored points), with the average shown (black points with line) and error bars showing the standard error of the mean. Identity of clam donors is listed in Table S1.

A four-hour incubation was chosen for these experiments as we found that proliferation of bacteria, protists, and unknown ciliates led to inconsistent cell survival in ASW beyond short-term incubation (consistent with Sunila et al). We recently found, however, that with the use of penicillin/streptomycin and notably, the addition of voriconazole, contaminant overgrowth could be controlled. We were therefore able to follow MarBTN cell survival long-term. We found that cells were able to survive far longer than four hours in 1× ASW, approximating typical marine conditions (Figure 2). We observed some variability in survival times for cells from different donor animals, but overall, we found that cells consistently survived longer at colder temperatures. At temperatures from 4°C to 16°C, an average of >48% of BTN cells survived for one week, and, at 4°C, 49% of cells survived for two weeks. For cells from one animal, >50% of BTN cells were still alive after one month at 4°C (living cells could still be detected after more than 8 weeks). This dramatically increases the amount of time BTN cells are known to survive in ASW, showing that BTN cells survive long enough to broadly disseminate through seawater to infect other clams.

**Figure 2.**
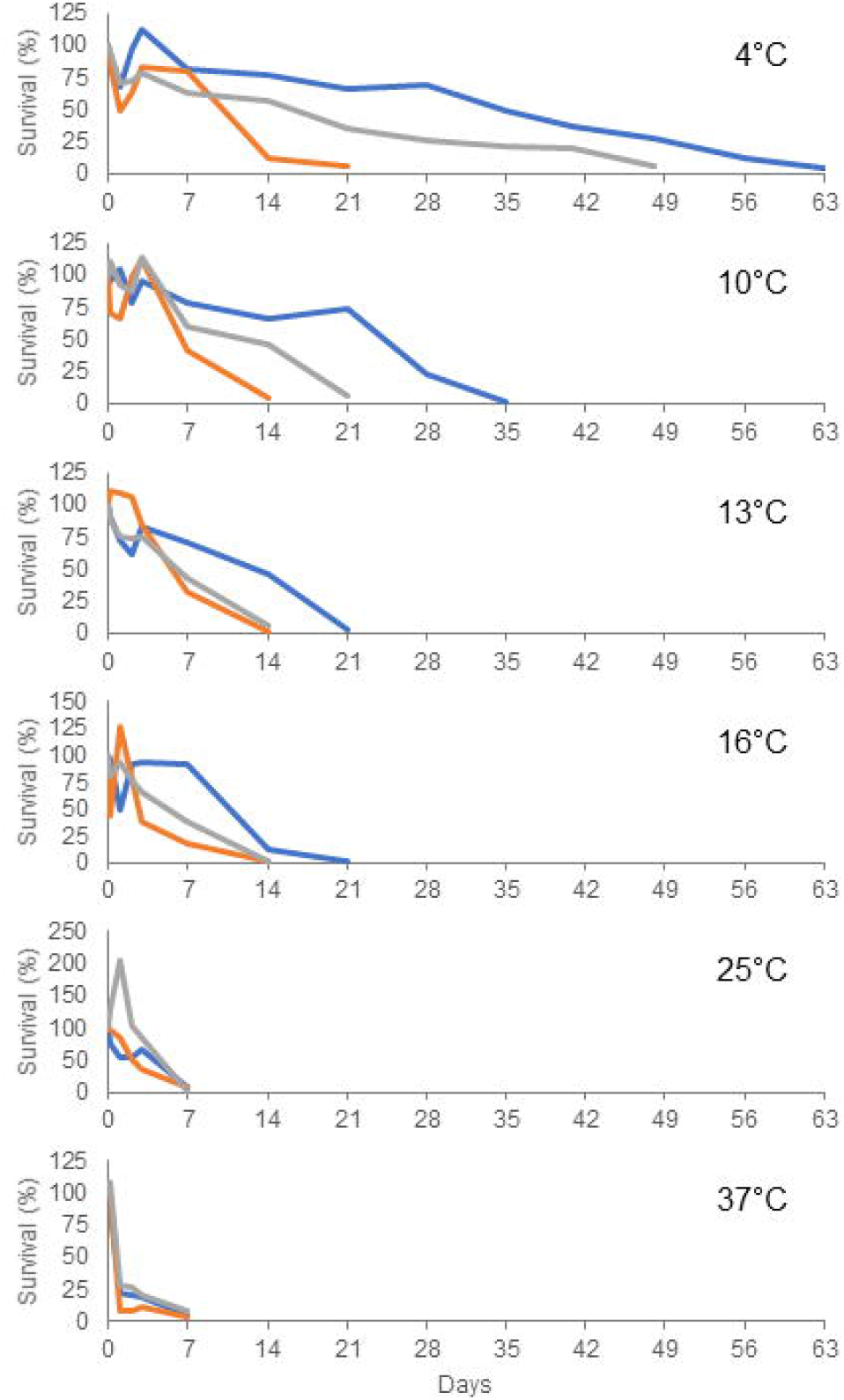
The effect of temperature on long-term survival of MarBTN cells in artificial seawater. BTN cells were collected from three different diseased clams and incubated in ASW (1.023 sg, pH 7.93, with penicillin/streptomycin/voriconazole). For each clam, cells were incubated in 4°C, 10°C, 13°C, 16°C, 25°C, and 37°C. Cell survival was monitored by resuspension and removal of an aliquot of cells, counted using erythrosine B, at 4 hrs, 1 day, 2 days, 3 days, 1 week, and weekly beyond that. Experiments were stopped when survival dropped below 10%.

For BTN to be spread through the water, cells need to survive, but they also need to get into the water from a diseased animal. To test whether BTN cells are released by diseased clams, we used qPCR of eDNA collected from both aquaria and natural water columns in regions with endemic BTN to look for two markers found only in *M. arenaria* BTN cells. Both cancer-specific primer pairs amplify specific integration sites of the LTR-retrotransposon *Steamer,* found only in BTN cells (*Steamer* is highly amplified within *M. arenaria* BTN cells, (23)). The cancer-HL03 primers amplify an insertion site cloned previously (23), and the *cancer-N1N2* primers amplify an insertion found near an ORF with high similarity to the gene *N1N2* (identified through preliminary analysis of MarBTN genome sequencing). As a control, primers in the conserved ORF of *N1N2* are used to amplify total copies of the *N1N2* locus. Both alleles in healthy cells amplify only with the healthy N1N2 primers, while MarBTN cells contain both the normal and the cancer-associated alleles and amplify with both primers (Figure 3A-C).

**Figure 3.**
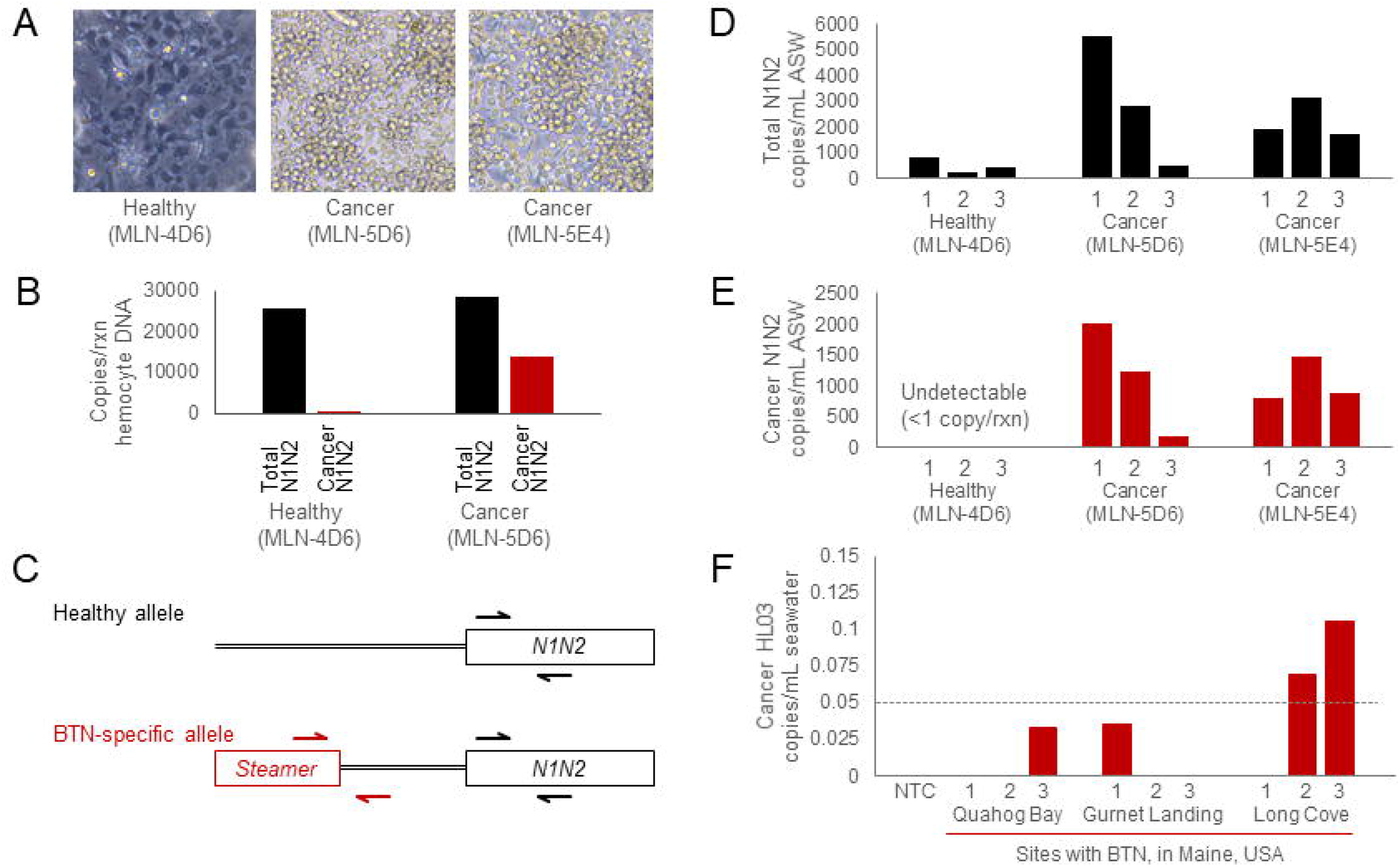
Detection of MarBTN eDNA from seawater in aquaria and from sites of known endemic BTN. Representative healthy and cancerous soft-shell clams were identified (***A***) through a screen of hemocyte morphology, and (***B***) the diagnosis was confirmed using a qPCR analysis of genomic DNA from hemocytes obtained from one of the diseased animal and the healthy animal used in the subsequent eDNA experiment. (***C***) The schematic shows the healthy *N1N2* allele and the cancer-associated allele, with arrows indicating the locations of the control primers (Total *N1N2*, black), used to determine the total number of clam alleles, and primers specific for the clam BTN lineage (Cancer *N1N2,* red), used to quantify BTN DNA. For eDNA analysis (***D-E***), each animal was housed in a separate aquarium, and eDNA was extracted from aquaria water on 3 sequential days. (***F***)Samples of water from sites in Maine with soft-shell clams known to have BTN were collected and eDNA was extracted. For each site, one water sample was collected and three sub-samples were extracted separately. qPCR analysis of the MarBTN-specific marker (Cancer-HL03) confirms detection of BTN DNA in the water. Copy numbers per μL DNA were converted to copies/mL, based on normalization to the total volume of water extracted. The dotted line shows 1 copy/reaction. For all qPCR, each sample was run in three reactions, and the values presented here are averages of the triplicate results. The average value was shown to be above zero only if the product was detectible in all triplicate reactions. Water was used as the no template control (NTC) and was undetectable in all three wells.

To test whether BTN-specific DNA can be released and subsequently detected in seawater, we housed one healthy and two highly neoplastic animals in separate, individual aquaria, changed the water, and then after 24 hrs collected a water sample for eDNA extraction. This was done for 3 consecutive days for each animal (Figure 3D-E). The qPCR data confirm that the healthy animal releases some normal DNA and no detectible BTN-specific DNA, while the heavily diseased animals release significant amounts of cancer-specific DNA across all three days, although the amount does vary from one 24-hour period to the next. This pattern is confirmed using the secondary HL03 cancer-specific primer set (Figure S1). Additionally, we can see that the ratio of cancer-*N1N2* to total-*N1N2* is between 0.4-0.5, suggesting that the majority of DNA in the water from both diseased animals came from released BTN cells (the cancer cells contain both an allele with the insertion and one without, so pure BTN cells have a 0.5 cancer-allele fraction).

The natural clam environment is far larger than a 1 L tank, so we next wished to test whether MarBTN-specific DNA could be found in wild environments where clam populations are known to be affected by endemic BTN. We chose three populations in Maine, collected surface water samples from each site, extracted eDNA, and performed qPCR (Figure 3F). Due to the potential for contamination with the control plasmid, only the HL03 marker was used for analysis of field samples, as this plasmid was not present in the lab where eDNA was extracted. These results showed levels far lower than aquaria levels with known heavily diseased animals, as expected, but we did observe MarBTN-specific amplification in two of the three sub-samples from Long Cove at a level above 1 copy/reaction (with amplification observed in all triplicate reactions for those two eDNA subsamples). This shows that BTN-specific DNA can be found in field samples of seawater in addition to being found in more concentrated laboratory conditions, again providing evidence for the hypothesized seawater-transmission of BTN.

## DISCUSSION

This study has shown that MarBTN cells can survive for many weeks in seawater under the right conditions, that they are acutely impacted by salinity but not pH, and that they can survive longer at colder environmental temperatures. We also show that eDNA from MarBTN cells can be detected in both aquaria and field samples, providing evidence for release of BTN cells from diseased animals for the first time. The previously proposed mechanism of transmission of BTN through seawater requires both long-term cell survival in the environment and release of BTN cells into the environment by diseased animals. This study provides evidence supporting both of those requirements.

This study largely agrees with the findings of Sunila et al. (10), showing the strong effect of salinity, but minimal effect of pH, and minimal effect of temperature on short-term survival except for toxicity at high temperatures. However, the cancer cells in the previous study demonstrated optimal survival in 10-15 ppt (approximately 1.0075-1.011 sg), a salinity level that was highly toxic to the cells in this study. Notably, the samples from that study were collected from northern Chesapeake Bay. In this estuary environment the surface seawater was 10 ppt, whereas the samples in the current study were taken from the coast of Maine, where the seawater has a much higher salinity. Sunila et al. had hypothesized an infectious cause for BTN, but it had not been confirmed at the time of that study, so it is unclear whether the cancer cells in that study were from the same lineage that is currently affecting New England and Prince Edward Island clams. The differences between these two findings strongly suggests that there has been evolution of BTN to survive in the seawater in which it must survive in order to transmit. This could represent evolution of two separate lineages within different environmental conditions, or it could represent divergence of a single lineage to better survive in marine vs. estuarine environments. Regardless, both studies showed that cancer cells were acutely sensitive to salinities lower than 10 ppt (1.0075 sg). To date, no DN has been observed in freshwater environments, so low-salinity environments may provide a potential “safe harbor” for bivalves, where transmissible cancer cannot survive.

We found clear evidence that BTN cells survive longer in the environment in colder temperatures, which may have implications for understanding the seasonality of BTN. BTN in soft-shell clams and other species have been reported to have seasonal fluctuations in prevalence (20, 24, 25), and these results suggest that transmission may be more likely in colder seasons, although there are additional unknown factors, such as the effect of temperature on the progression of disease.

A very recent study of the MtrBTN2 lineage of transmissible cancer, known to infect four *Mytilus* species around the world (14–16), has shown that these cancer cells also can survive for a few days in seawater (26). The authors assayed cell survival at 18°C; our results showing longer survival of MarBTN at lower temperatures suggest that their finding of 6-day survival may be an underestimate. It will be interesting to determine in the future whether our finding of the effects of salinity and temperature on survival are the same across BTNs from different species.

One limitation of our study is that detection of MarBTN-specific eDNA does not confirm that live cells are in the environment, only that the DNA can be detected. However, given the fact that BTN cells in different animals are identical to each other, and that BTN cells can survive well in the marine environment, it seems reasonable to conclude that eDNA is detecting live cells. This study provides the proof of principle for an eDNA assay that can be used to determine the timing of cell release during BTN progression. It can also be used to identify the presence of BTN in field samples, potentially serving as a non-invasive proxy for monitoring disease in the wild and possibly reducing the requirement for more invasive and expensive screening of animals for disease.

In this study, we show evidence supporting the long-term survival of MarBTN cells and release of MarBTN cells from diseased animals. Overall, these data provide proof of principle supporting the transmission of BTN through the seawater as a pathogen, and they establish new methods to investigate the mechanisms of BTN survival, progression, and spread.

## ACKNOWLEDGEMENTS

We acknowledge the assistance of David Wilson for collection of seawater samples in Maine and Bret Pearson (University of Toronto) for suggesting the use of voriconazole.

## SUPPLEMENTARY MATERIALS

Table S1. Collection information of soft-shell clams (*Mya arenaria*) used in this study

Table S2. Primers used in qPCR and cloning

Figure S1. Detection of MarBTN eDNA from seawater in aquaria with secondary qPCR marker

