## Supplementary material for "Survival and detection of bivalve transmissible neoplasia from the soft-shell clam *Mya arenaria* (MarBTN) in seawater": Table S1 and S2

Table S1. Collection information of soft-shell clams (*Mya arenaria*) used in this study

| Animal ID | Cancer in hemolymph[1] | Arrival date | Harvest location | Figure (color) |
|---|---|---|---|---|
| FFM-4D7 | 100% | 10/22/2019 | Lubec, ME | 1A (orange) |
| FFM-4G8 | 100% | 10/22/2019 | Lubec, ME | 1A (blue) |
| FFM-6B8 | 75% | 10/31/2019 | New Brunswick, ME | 1A (grey) |
| FFM-26C1 | 100% | 10/1/2021 | Friendship, ME | 1B & 2 (blue) |
| FFM-26C9 | 100% | 10/7/2021 | Friendship, ME | 1B (orange) |
| FFM-26D8 | 100% | 10/7/2021 | Friendship, ME | 1B & 2 (grey) |
| FFM-5D4 | 75% | 10/31/2019 | New Brunswick, ME | 1C (light blue) |
| FFM-7A12 | 100% | 12/10/2019 | Jonesport, ME | 1C (green) |
| FFM-7F4 | 100% | 12/10/2019 | Jonesport, ME | 1C (dark blue) |
| FFM-8C3 | 100% | 1/7/2020 | Brunswick, ME | 1C (orange) |
| FFM-9A1 | 100% | 1/7/2020 | Brunswick, ME | 1C (blue) |
| FFM-9A12 | 100% | 1/7/2020 | Brunswick, ME | 1C (grey) |
| FFM-26C10 | 100% | 10/7/2021 | Friendship, ME | 2 (orange) |
| MLN-4D6 | 0% | 1/29/2019 | ME | 3, S1 |
| MLN-5D6 | 100% | 2/19/2019 | ME | 3, S1 |
| MLN-5E4 | 100% | 2/19/2019 | ME | 3, S1 |

1 Cancer in hemolymph was first estimated through microscopic analysis of hemolymph. Only highly diseased animals were selected for use as donors for BTN cell survival experiments

Table S2. Primers used in qPCR and cloning

| qPCR target | Control plasmid | Primer name | Primer Sequence (5'-3') |
|---|---|---|---|
| Cancer-N1N2 | pCR-SteamerLTR-N1N2 | ClamLTR-F3<br>N1N2can-R3 | TTCAATCATTCAACGCATAACC<br>TCGCTGAGAATTTTTCGGTGT |
| Total-N1N2 | pCR-SteamerLTR-N1N2 | N1N2-F3<br>N1N2-R1 | CCCAGGGCAAGAGGAATATGGT<br>GGATACTGCAAGCTTCTTGGAA |
| Cancer-HL03 | pIMHL03c2-EF1α | ClamLTRF2<br>IMHLO3c2-R2 | ACATGCACATTAAAAGTTATCG<br>TCTGGGTCATGAATAACGTCA |
| EF1α | pIMHL03c2-EF1α | ClamEF1-F3<br>ClamEF1-R2 | GGGAAAAGAGGGCAAGGTGAC<br>TTTCTTCTTCCCACCGACTGC |
| Cloning primers | | | |
| | pCR-SteamerLTR-N1N2 | ClamLTR-F2<br>98171_conR1 | ACATGCACATTAAAAGTTATCG<br>GGATACTGCAAGCTTCTTGGAA |
| | pIMHL03c2-EF1α | ClamLTR-F2[1]<br>ClamLTR-R1[1]<br>ClamA-EF1aFor[2]<br>ClamNS-EF1aRev[2] | ACATGCACATTAAAAGTTATCG<br>TTAGTATAGCCAATACTGTTAC<br>tagggcccGAAGGATGAGGGAAAAGAGGG<br>atGCGGCCGCatcctgcaggCACCTTTTCCTGCTATGGTGC |

1 Cloning of the *Steamer* fragment to generate pIMHL03c2 was done via inverse PCR, as described in Arriagada et al. 2014.

2 Primers were used to amplify *EF1α* from genomic DNA, the product was cut with ApaI and NotI (NEB), and ligated into pIMHL03c2.
