## Supplementary figures and images for "Survival and detection of bivalve transmissible neoplasia from the soft-shell clam *Mya arenaria* (MarBTN) in seawater"

### Figure S1

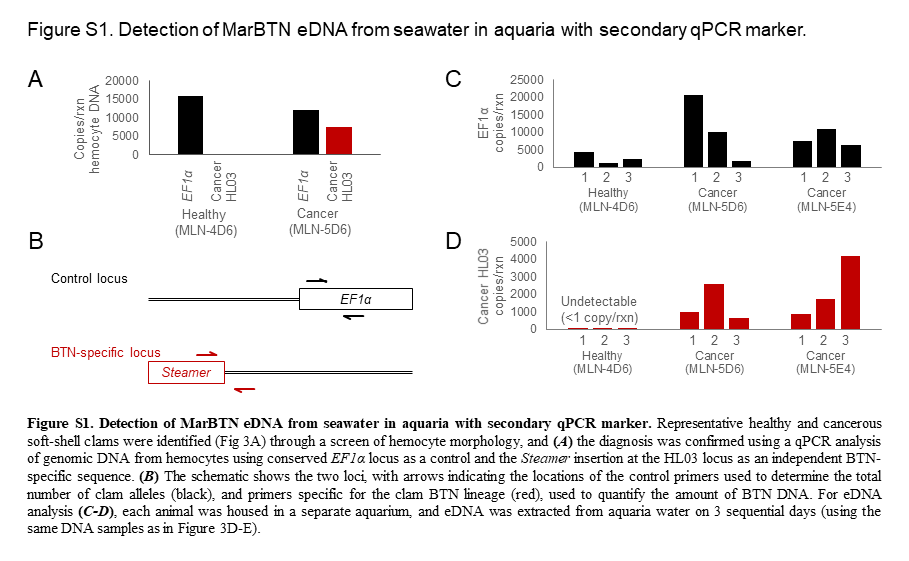
